# Cluster replicability in single-cell and single-nucleus atlases of the mouse brain

**DOI:** 10.1101/2025.02.24.639959

**Authors:** Leon French, Hamsini Suresh, Jesse Gillis

## Abstract

Single-cell RNA sequencing has advanced our understanding of cellular heterogeneity. Ensuring the replicability of identified cell clusters across studies is essential for determining their biological robustness. We assess the replicability of cell clusters identified in two large mouse brain atlases, one generated using single-cell RNA sequencing and the other with single nuclei. Both profile over 4 million cells and group them into over 5000 clusters. Using transcriptome-wide neighbor voting, we identify 2009 reciprocally matched cluster pairs with consistent spatial localization and coordinated gene expression, which were also observed in datasets from multiple species. Reciprocal clusters are enriched in the cerebellum, where lower diversity aids replicability, while the hypothalamus’s heterogeneity limits agreement. Distinguishing close clusters is much more challenging than differentiating a cluster from most others, especially when using marker genes. By incorporating replicability data, we provide a stronger foundation for investigating the newly identified clusters and their biological meaning.

## Main

The advent of single-cell RNA sequencing has significantly revolutionized our understanding of cellular diversity in the brain ^1,2^. Large-scale transcriptomic atlases provide invaluable resources for dissecting cellular heterogeneity ^3^. Validating atlas consistency is vital for their practical application in neuroscience research ^4^. Two recent mouse brain atlases, each independently identifying over 5,000 cell clusters using single-cell and nucleus sequencing, underscore the magnitude of this challenge ^5,6^. Assessing cluster replicability, or the degree to which cell clusters identified in one study are consistent with those found in another, is critical to any claim that they represent reproducible biology, such as cell types ^7^.

Prior work by the BRAIN Initiative Cell Census Network (BICCN) has identified 70 highly replicable cell types in the mouse primary motor cortex ^8^. This analysis leveraged seven transcriptomic datasets that differed in preparation methods (single nuclei vs. whole cells) and sequencing platforms (10x Genomics v2, v3, and SMART-Seq). Notably, the authors observed that replicability decreased as cells were subdivided into finer partitions. While single-nucleus sequencing detects fewer transcripts per cell, previous investigations have found it to possess comparable sensitivity to single-cell sequencing for identifying cell types in brain tissue ^9–11^. However, these comparisons were limited to fewer than 12 coarse cell types, and single-nucleus sequencing may not be sufficient to detect disease-associated cell states ^12^. From an evolutionary perspective, six retinal cell classes were shown to be highly conserved across 17 species, with transcriptomic similarity decreasing as evolutionary distance increased ^13^. These observations raise the following questions: Do these findings extend to the thousands of cell clusters identified in the two recently published whole-brain studies? If so, what are the properties of these replicable clusters?

This study deeply characterizes the replicability of cell clusters identified in the two central BICAN reference large-scale mouse brain atlases. We systematically evaluate cluster similarity through marker gene analysis and an expression learning formalism using transcriptome-wide neighbor voting between all cells. We further characterize the replicable clusters, focusing on their regional enrichment, spatial alignment, coordinated gene expression, and representation across diverse mouse and cross-species brain datasets. By rigorously assessing cluster agreement, we aim to highlight both the limitations and strengths of large-scale transcriptomic atlases in capturing cellular heterogeneity as well as providing a firm foundation for subsequent work.

## Results

### Basic properties of the single-cell mouse brain transcriptomic atlases

We focused on the recently published single-cell and single-nucleus atlases that transcriptomicly profiled the entire mouse brain ^5,6^. The single-cell atlas contains transcriptomic profiles of over 4 million cells grouped into 5,322 clusters. Similarly, the single-nucleus atlas assayed expression in 4.4 million nuclei that have been grouped into 5,030 clusters (Fig. 1a). The broad regional source of the cells varies with more cells sampled from the thalamus, midbrain, brainstem, and cerebellum in the single-nucleus dataset while the single-cell atlas captured more cells in the hypothalamus and cerebral cortex (Fig. 1b). Similar differences were observed when comparing the number of clusters detected per dissection region (Fig. 1b). The number of clusters appearing within the dissection regions showed a higher correlation (Spearman rho = 0.91) between the two studies compared to cell counts (Spearman rho = 0.71). This suggests agreement in the degree of cellular heterogeneity across broad regions despite sampling differences. The cluster sizes in both atlases range from 9 to 264,669 cells in the single-cell dataset and 15 to 552,018 in the single-nucleus dataset. Consequently, a small fraction of the clusters contain most of the cells with a higher concentration of nuclei compared to the cells (Fig. 1c). For example, the ten largest clusters contain 28.9% and 52.4% of the cells in the single-cell and single-nucleus datasets, respectively. Overall, while both atlases provide extensive coverage of the mouse brain, they exhibit differences in cell sampling and cluster distribution, with a significant portion of cells concentrated in a few large clusters.

**Figure 1:**
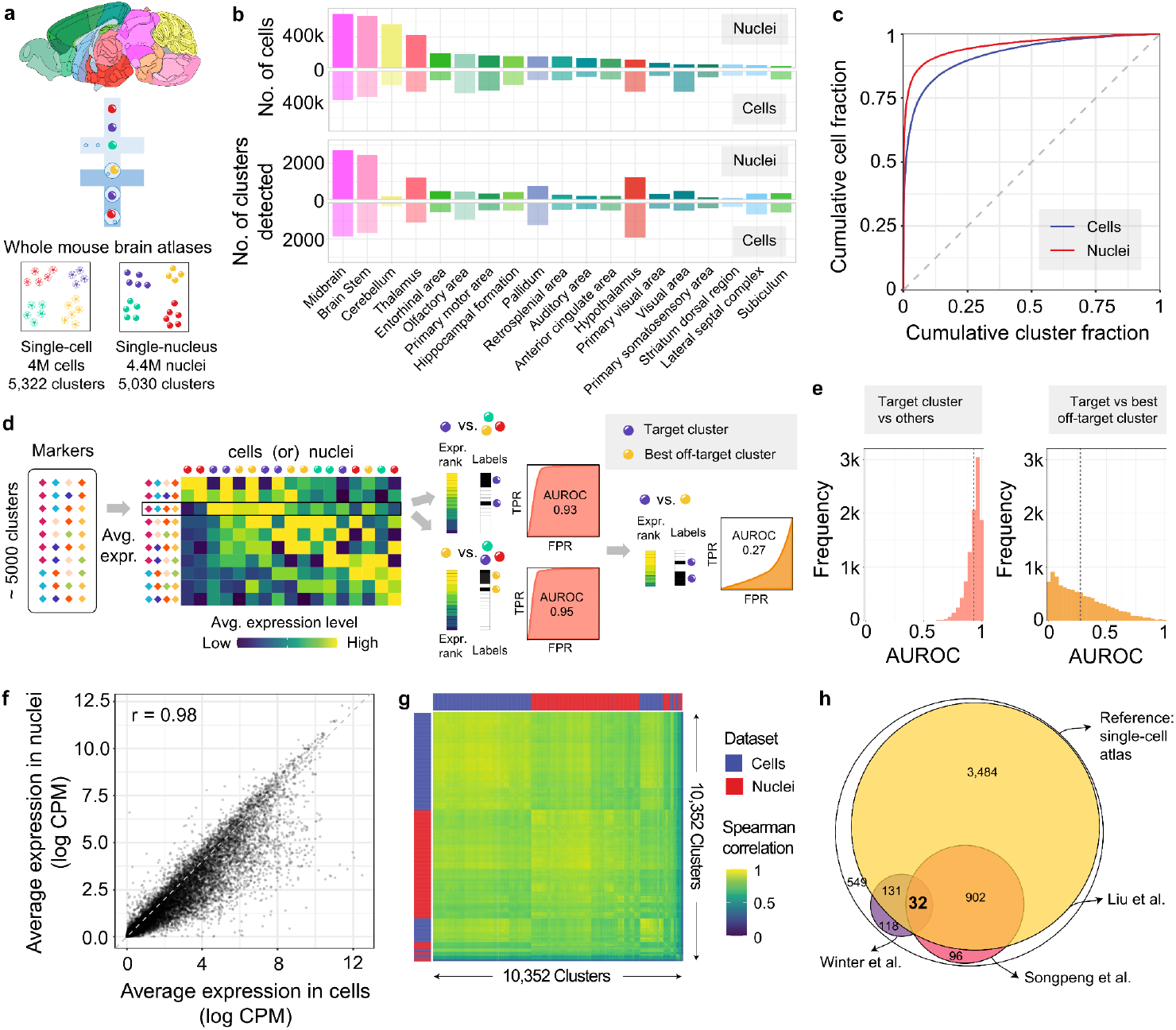
Markers are insufficient for aligning single-cell and single-nucleus whole-brain atlases. **a**, Summary of the two whole brain mouse atlases. **b**, Bar plots showing the distribution of cells and detected clusters across dissection regions in the atlases. **c**, Cumulative proportions of cells and clusters ranked by size, shown for single-cell (blue) and single-nucleus (red) clusters. **d**, Schematic illustrating how author-provided markers are tested for detection of their target cluster cells relative to all clusters and similar clusters within an atlas. **e**, Distribution of AUROC values for detecting target clusters relative to all clusters (left, red) and most similar non-target clusters (right, orange). Mean AUROC values are marked with dotted lines. **f**, Scatter plot showing average gene expression (log CPM) in cells versus nuclei when averaged across cluster centroids. Each dot represents a gene, showing a strong correlation (Spearman’s r = 0.98). **g**, Heatmap of genome-wide expression profile correlations between cluster centroids from both atlases. **h**, Euler diagram depicting the overlap between clusters identified by Liu et al. (yellow), Winter et al. (purple), and SongPeng et al. (pink) when mapped to the single-cell atlas.

### While effective in the whole brain context, marker genes do not separate similar clusters

Since marker genes are commonly used to annotate cell types, we evaluated the specificity and recurrence of the author-provided marker lists before examining genome-wide expression profiles of the atlases. The median number of markers per cluster was four for both atlases, with 3,853 unique genes listed as markers. For a given target cluster and its markers, we ranked every cell by the average z-scored expression of the markers and tested if the target cluster cells rank higher than other cells using the area under the receiver operating curve statistic (AUROC) (Fig. 1d). This statistic can be interpreted as the probability that the average gene expression of the markers will rank a cell of the target cluster higher than any other cell. As expected, the markers rank the target cells highly with mean AUROC values of 0.945 and 0.926, for the single-cell and single-nucleus datasets, respectively (Fig. 1e). When compared to other clusters, the performance is weaker with only 304 single-cell and 287 nuclei target clusters having the highest AUROC value when all clusters are tested. In other words, marker sets were often more specific to an off-target cluster. Furthermore, the top ranked clusters collapse to less than 1,000 unique clusters when all marker sets are tested within an atlas. To detail this more directly we constrained the set of tested cells to those of the target cluster and the best off-target cluster to determine if the markers could discriminate between similar clusters (Fig. 1d). In this local setting, cells from target clusters are ranked below the best off-target cluster on average (mean AUROC=0.27 to 0.28, Fig 1e). Langlieb et al. acknowledged the difficulty in finding markers that can effectively discriminate between very similar clusters and, therefore, provided markers that also identify the cluster neighbours ^6^. However, for our analyses, we used only their markers that were obtained without this relaxed threshold. Overall, these findings underscore the challenge of using a few markers for cluster identification.

Having characterized the specificity of markers within the individual atlases, we next compared the similarity of clusters across datasets by testing the overlap of the marker gene lists. If the same clusters are found in both atlases, then their markers should overlap. After correcting for the 23 million pairwise cluster comparisons, 30 pairs of clusters have a statistically significant overlap of markers (hypergeometric test, Bonferroni correction). A relaxed threshold that corrects for the cluster count in each dataset (∼5,000), identifies 1,826 pairs of clusters with overlapping markers. This limited agreement between marker genes suggests that genome-wide expression profiles are better suited to examine cluster similarity.

To broadly gauge gene expression similarity between the atlases, we next compared average expression at different levels. Across the genome, when averaged across cluster centroids, the correlation between single-cell and single-nucleus atlases is high (Spearman’s rho = 0.98, Fig. 1f). As expected, some genes are expressed more in whole cells than in nuclei due to the loss of cytoplasmic transcripts. Correlating average genome-wide expression profiles across clusters shows a clear grouping within each atlas (Fig. 1g). Though there is strong per-gene correlation in average expression at the atlas level, the differences between clusters are not strong enough to override atlas-specific signals.

Three companion studies from the Brain Initiative Cell Census Network 2.0 have aligned their single-cell clusters to the single-cell atlas of the mouse brain using advanced integration approaches. Liu et al. mapped 87.7% of the single-cell atlas clusters to at least one of their 4,673 cluster-by-spatial cell groups, which were derived from single-cell methylomes. However, most of these mappings were ambiguous between the two datasets, with only 145 one-to-one cluster pairs identified. In a second study, which examined chromatin accessibility across the mouse brain, 19.5% of the single-cell atlas clusters were mapped to 1,482 distinct brain cell populations from the snATAC–seq data (Songpeng et al.). After integration, the median proportion of cells annotated to the best-matched single-cell atlas clusters was 20.7%. A third transcriptomic study focused on spinal projecting neurons from eight regions (Winter et al.). It found that the 76 types of projecting neurons mapped to 291 of the 5,322 single-cell atlas clusters. This mapping was also ambiguous, with no one-to-one mappings. By intersecting the mapped sets of clusters from these three studies, we tested the replicability of specific single-cell atlas clusters. Only 32 clusters were consistently mapped across all three studies, fewer than expected by chance (expected: 48.7, empirical p < 0.01, Fig. 1h). A larger overlap appeared when excluding the projection study, which focused on neurons. Specifically, 934 scRNA-seq clusters replicated in the brain-wide epigenetic studies (expected = 889, p < 0.001). The limited and ambiguous mappings from advanced integration methods motivated our comparison of the single-cell and single-nucleus transcriptomic mouse brain atlases.

### MetaNeighbor reveals cross-atlas cluster relationships

Guided by the limits of marker genes and previous integration results, we used MetaNeighbor to assess replicability of the clusters in the two atlases ^14^. As diagrammed in Fig. 2a, MetaNeighbor builds a cell-cell similarity network weighted by the Spearman correlation across a set of highly variable genes (HVG). Cross-dataset neighbor voting based on this network is then used to provide interpretable cluster similarities (AUROC scores). Effectively, each cell is asked how well it matches to all cells in the other dataset and this is compared with other cells within the same dataset, correcting for global data shifts. Importantly, MetaNeighbor has been optimized for large single-cell datasets ^15^. The resulting AUROC heatmap resulting from MetaNeighbor when ran on all 8.3 millions cells, 3,534 highly variable genes, and 10,352 clusters shows a large block of very similar neuronal clusters. After expanding contrast for visualization of high AUROC values, a set of diagonal blocks of replicable clusters from both datasets is evident (Fig. 2b). However, at this scale, this heatmap does not capture the differences between very similar clusters because each pixel represents the AUROC values of several clusters due to resolution limits. This is clear in an expanded view of the 246 non-neuronal clusters (Fig. 2b). When zoomed in, clear blocks of similar glial clusters that cross datasets are grouped with smaller blocks that indicate hierarchical structure. While similar clusters are apparent in this global comparison, focused analysis is needed to evaluate highly concordant clusters.

**Figure 2:**
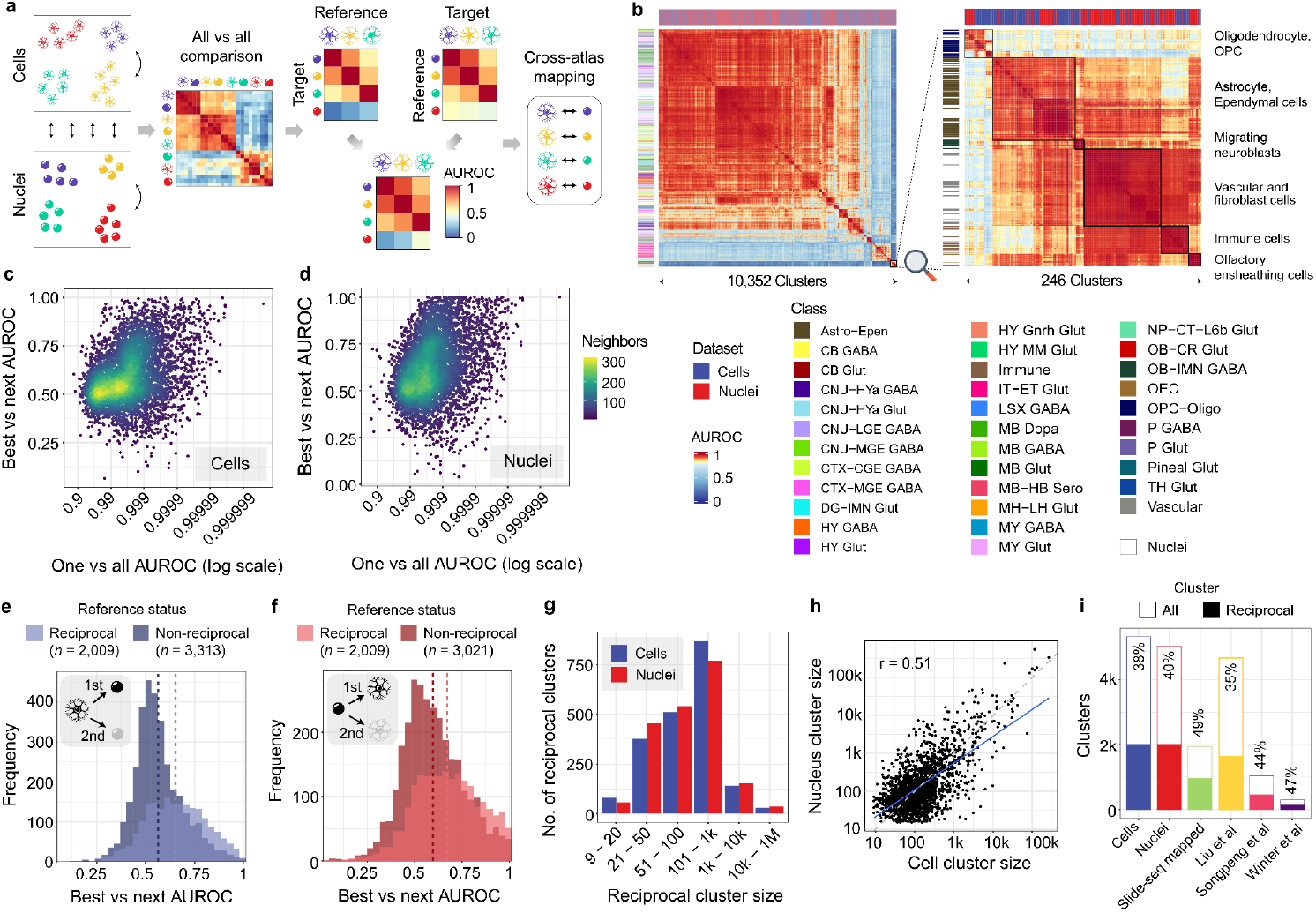
Identification of reciprocally best-matched cell type clusters between the two whole brain atlases using MetaNeighbor. **a**, Schematic showing the unsupervised MetaNeighbor framework used to select replicable clusters. Cell type-specific expression profiles learned from a reference dataset (single-cell) are used to predict similar cells in the target dataset (single-nucleus) and vice versa. Reciprocal top hits between the two datasets represent a set of highly replicating cell clusters between the atlases despite the compositional and technical variability in the underlying data. **b**, Heatmap showing ‘one_vs_all’ MetaNeighbor scores for 10,352 clusters (left), and for 246 select non-neuronal clusters (right) obtained by zooming into the bottom right corner of the complete heatmap. Clusters are labeled by dataset (top) and class for the single-cell clusters (left). Transcriptional similarity between pairs of clusters is depicted using a color scale that ranges from yellow to red for a narrow range of high replicability scores (0.8 to 1 AUROC) for better contrast. **c-d**, Scatterplots showing the relationship between global and local AUROC scores, using the single-cell (left) and nuclei (right) clusters as the references. The x-axis displays global scores (best one-vs-all AUROC) on a logarithmic scale, where 1 represents perfect identification. The y-axis shows local scores (best-vs-next AUROC) for the same reference clusters. Points are colored based on the density of neighboring points. **e-f**, Histograms indicating the best-vs-next AUROC scores for non-reciprocal (darker shade) and reciprocal top hit (brighter shade) clusters from the perspective of the cell (blue) and nuclei (red) based reference datasets. **g**, Barplots showing the number of reciprocal top hit clusters across different size bins in the single-cell (blue) and single-nucleus (red) atlases. **h**, Scatter plot comparing the log scaled sizes of reciprocal top hit pairs between the two atlases (linear regression fit shown as blue line, grey dashed line represents the line of equality, and Spearman correlation coefficient at top left). **i**. Bar plots marking the count of total clusters (white bar) and reciprocal top hit clusters in each dataset (colored bars).

We next summarize the full AUROC matrix by examining the best cross-dataset AUROC scores for each cluster. Almost all clusters identify another in the other atlas with an AUROC value above 0.95. Mirroring the marker set results, these high AUROC scores which are relative to all cells are not repeated when close clusters are compared. This is measured using the “best-vs-next” AUROC score, which tests, for a given reference cluster, if cells from the two most similar clusters in the other atlas can be discriminated based on expression similarity to the cells from the reference cluster. The average of this local AUROC score is 0.598 for single-cell and 0.627 for the single-nucleus reference clusters (Figs. 2c-d).

Between the two atlases, we identified 2,009 cluster pairs as reciprocal top hits, meaning they preferentially have the best AUROC scores with each other (Supplementary Table 1). When compared to their most similar clusters, these pairs have higher average “best-vs-next” AUROC scores, exceeding 0.65 (Figs. 2e-f). Similarly, selecting markers that distinguish both sides of the reciprocal pairs yielded fewer than 6% of clusters that could be locally discriminated with an AUROC greater than 0.95 using four marker genes (Supplementary Note 1 and Supplementary Figure 1). To refine the reciprocal pairs, we applied stricter thresholds to identify a high-confidence subset (Supplementary Table 8) and explored merging clusters to increase the number of matches. However, since these approaches either reduced coverage or yielded only marginally more reciprocal top hits, we focused on the full set of 2,009 pairs (Supplementary Note 2).

Reciprocal top hits are observed across the range of cluster sizes. Larger single-nucleus clusters are more likely to be reciprocal, with a median size of 95 cells compared to 70 cells for non-reciprocal hits. In contrast, smaller single-cell clusters are more likely to be reciprocal, with a median size of 106 cells versus 134 for non-reciprocal hits. In aggregate, the majority of profiled cells are found in reciprocal best hit clusters, with 76.2% of nuclei and 50.2% of cells in these clusters (Fig. 2g). Although MetaNeighbor’s AUROC scores are independent of cluster size, the similar sizes of paired reciprocal clusters suggest comparable sampling of these paired clusters (Spearman r = 0.51, Fig. 2h).

Analysis of the local context shows that larger reference clusters are more effective at distinguishing similar clusters in the other atlases. Specifically, reference clusters with more than 100 cells demonstrate better discriminatory power, with 36.4% having statistically significant ‘best-vs-next’ AUROC scores (Wilcoxon rank-sum test, Bonferroni correction, Supplementary Figure 2). In contrast, only 25.9% of the smaller clusters pass this threshold. Additionally, large reference clusters from the single-cell atlas demonstrate better cross-study discrimination ability compared to single-nucleus clusters. Overall, a substantial proportion of clusters participate in reciprocal best hits, which vary in size and demonstrate improved local discrimination ability.

We next intersected our reciprocal hits with the clusters found in separate integration efforts to test whether they were more likely to be observed. In the single-nucleus study, 49.2% of the 1,937 clusters detected in the companion spatial Slide-seq experiment were reciprocal top hits, exceeding the expected 40% (p < 0.0001, Fig. 2i). For the three studies that mapped to the single-cell atlas, the Liu et al. mappings were depleted for reciprocal hits (36%) while the Songpeng et al. (44.4%) and Winter et al. (54.3%) mappings were enriched (all p < 0.0001, Fig. 2i). These results indicate that while there is some variation, our reciprocal top hits are generally observed at or above the expected rates, reflecting moderate enrichment in independent integration efforts.

### Characterization of the reciprocal best hit cluster pairs

To better understand the 2,009 reciprocal best hits we examined their placement in the broader cell type taxonomies and their locations in the mouse brain. We first tested if the reciprocal clusters were enriched for specific classes from the Yao et al. single-cell based taxonomy. Of the 34 classes, those containing clusters of GABAergic neurons from the medulla, midbrain, and pons; glutamatergic neurons from the medulla and midbrain; and immature GABAergic neurons from the olfactory bulb were enriched for reciprocal hits (hypergeometric test, pFDR < 0.05, Supplementary Table 2). Across the meta-cluster classification from the single-nuclei dataset from Langlieb et al. (n=161 with ten or more clusters), astrocytes; and neurons annotated as ‘thalamic projection’, ‘brainstem’, ‘medullary’, ‘inhibitory olfactory’, ‘pontine and medullary’, ‘tegmental and pontine’, and ‘midbrain and pontine’ were enriched (Supplementary Table 3).

To directly assess the spatial agreement of cluster locations, we mapped dissection annotations from the atlases to 19 common regions. For the 2,009 reciprocal best hit pairs, the dissection region containing the highest proportion of cells matched between datasets for 80.8% of the pairs. For the 996 cluster pairs with cells from two or more dissection regions, the top two regions with the highest proportion of cells matched for 45.6% of the pairs. As a comparison, a set of 972 best-matching cluster pairs based on the single-cell atlas markers exhibited lower matching rates—31.6% for the top region and 13.6% for the top two regions.

To examine reciprocal top hit clusters at finer spatial resolution than dissection regions, we called each cell in the MERFISH dataset used by Yao et al. using the same method with the single nucleus cluster centroids (Fig 3a). For each cell, we tested if calls generated from the two separate atlases corresponded to a reciprocal best hit pair. For example, seven cells that map to both sides of the smallest reciprocal hit demonstrate spatial co-localization (Fig 3b). In this case, three cells are annotated to the single-cell based reciprocal cluster but could not be confidently called when using the single-nucleus data as a reference (average correlation score < 0.5). Across all 4.3 million cells, 3.9 and 2.0 million have been assigned to a cluster based on single-cell and single-nucleus references respectively. Of these, 768,245 cells are assigned to both sides of a reciprocal best hit pair. Visualization of these cells across four coronal sections (Fig. 3c) reveals a striking enrichment in the cerebellum and depletion in the middle cortical layers of the cerebral cortex. The cerebellar pattern is primarily driven by the most frequent cluster pair (Ex_Gabra6_Gabrd/cluster_ID:5201), which marks 151,723 granule cells. The second most abundant reciprocal cluster pair consists of 65,194 cells annotated as cerebellar GABAergic neurons (Inh_Tfap2a_Sla/cluster_ID:5188). Although these neurons are small, cell density does not appear to drive the enrichment, as gridded images that normalize for cell density show a similar spatial pattern (Fig. 3c, bottom).

**Figure 3:**
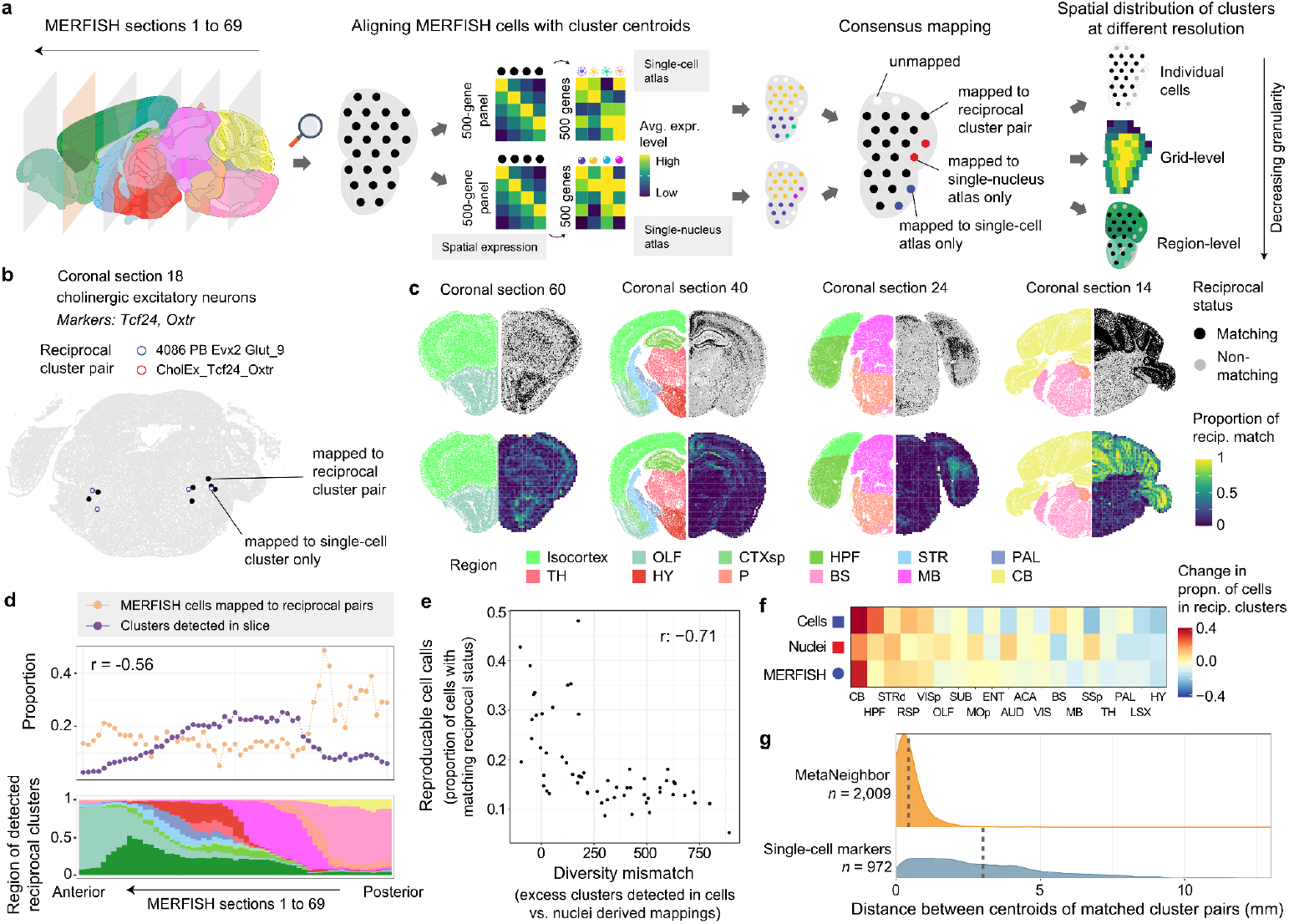
Distribution of reciprocally matched clusters in a spatial transcriptomic atlas of the whole brain. **a**, Schematic illustrating the workflow used to align individual cells in MERFISH sections to clusters in the single-cell and single-nucleus atlases. **b**, MERFISH coronal section (coronal slice 18) highlighting 7 cells that are mapped to both clusters of an example reciprocally matched pair. Additionally, four cells mapped only to the single-cell side of the cluster pair are marked in blue. **c**, Representative MERFISH sections showing the spatial distribution of cells mapped to reciprocally matched cluster pairs at single-cell (top) and coarse grid-level (bottom) resolutions. Each section displays cells’ spatial locations colored by broad CCFv3 regions (left hemisphere), and the distribution(top)/proportion(bottom) of cells mapped to reciprocally matched cluster pairs (right hemisphere). **d**, Line plot depicting the proportion of cells mapped to reciprocally matched cluster pairs (orange dots) and detected clusters (purple dots) within coronal sections ordered from anterior to posterior along the x-axis. Stacked bar chart (bottom) showing the distribution of the detected reciprocally matched clusters across the same slices, colored by broad CCFv3 regions. **e**, Scatterplot showing the relationship between reproducible cell calls (y-axis) and the mismatch in cluster diversity (x-axis) for each slice. Reproducible cell calls are defined as the proportion of cells mapped to reciprocal pairs. The diversity mismatch represents the difference in the number of clusters detected using single-cell versus single-nucleus references. The Spearman correlation between these measures across the slices is -0.71 (top right). **f**, Heatmap indicating the relative enrichment of cells in reciprocally matched clusters within aligned regions from the single-cell, single-nucleus, and MERFISH spatial atlas data. Columns (regions) are ordered from highest average enrichment (left) to depletion (right). **g**, Density plots of the distances in mm between centroids of Metaneighbor reciprocal top hits (top) and marker-derived (bottom) cluster pairs in the MERFISH dataset. Mean centroid distances are indicated by dashed lines. Anatomical template image is from the Allen Mouse Brain Reference Atlas ^16^.

Across the coronal slices, the most anterior and posterior slices exhibit higher proportions of cells assigned to reciprocal best hit clusters (Fig. 3d). This proportion is inversely correlated with the diversity of clusters detected in each slice (Spearman r = -0.56, Fig. 3d). To further investigate this relationship, we downsampled the number of MERFISH cells identified using the single-cell reference to match the number of nuclei-based calls. We then calculated the difference in detected cluster counts between between the two references. Across the slices, the excess detection of single-cell clusters in this diversity mismatch measure is negatively correlated with the proportion of reciprocal best hits (Spearman r = -0.71, Fig. 3e). This association suggests that differences in sampling depth and clustering resolution can influence cluster replicability between the two atlases.

Across 19 aligned dissection regions, we find that the proportion of cells mapped to reciprocal best clusters is consistent with the finer-resolution MERFISH results. Specifically, cells in the cerebellum and the hippocampal formation are more likely to be mapped to reciprocal best clusters, while the hypothalamus and the nearby lateral septal complex are depleted for cells from reciprocal clusters (Fig. 3f).

### Spatial proximity of reciprocally matched clusters

To test similarity between reciprocally matched clusters in the spatial domain, we calculated the distances between their centroids. The average distance between the centroids of a reciprocal best hit cluster pair is 0.428 mm. In contrast, for the 972 best-matching cluster pairs derived from markers, the average distance was significantly larger at 3.12 mm (Fig 3g). Within the 2,009 reciprocal top-hit pairs, the next closest clusters from the MetaNeighbour analyses are also farther apart, with a mean distance of 0.95 mm (Wilcoxon rank-sum test, p < 0.0001). The close spatial proximity of these reciprocally matched clusters is notable, as spatial information was not used during the clustering or matching processes, making this convergence especially reassuring.

### Coordinated gene expression across the matched clusters

To provide a genome-wide comparison of the reciprocally matched clusters, we compared their per-gene expression profiles across clusters. We measured coordinated expression across the paired cluster expression centroids (Fig. 4a). For this analysis, we excluded the 3,534 genes that were used to determine the reciprocal best hit pairs. For the remaining 17,450 genes, the per-gene average Spearman correlation across the 2,009 reciprocal clusters was 0.357 (Fig. 4b). We randomly sampled the number of paired clusters to test if this association is sensitive to the cluster count. The average correlation remained stable across the range of sizes, suggesting that smaller atlases would provide similar levels of coordinated expression. To test specificity, we examined whether different gene pairs have stronger correlations than the same gene pairs between the two atlases. The average percentile ranking of same gene pairs was 94.9%, which slightly decreased as cluster pairs were downsampled (Fig 4c). For 34.8% of the genes, their coordinated expression was higher than all correlations between that gene and any other (Fig. 4d). This metric was the most sensitive to the number of clusters, dropping to 27% on average for 10% of the original clusters. Unsurprisingly, we find that genes involved in synapse-related functions were enriched for higher coordinated expression (Fig. 4e, Supplementary Table 4). This analysis highlights the robustness and specificity of coordinated gene expression in the reciprocally matched clusters.

**Figure 4:**
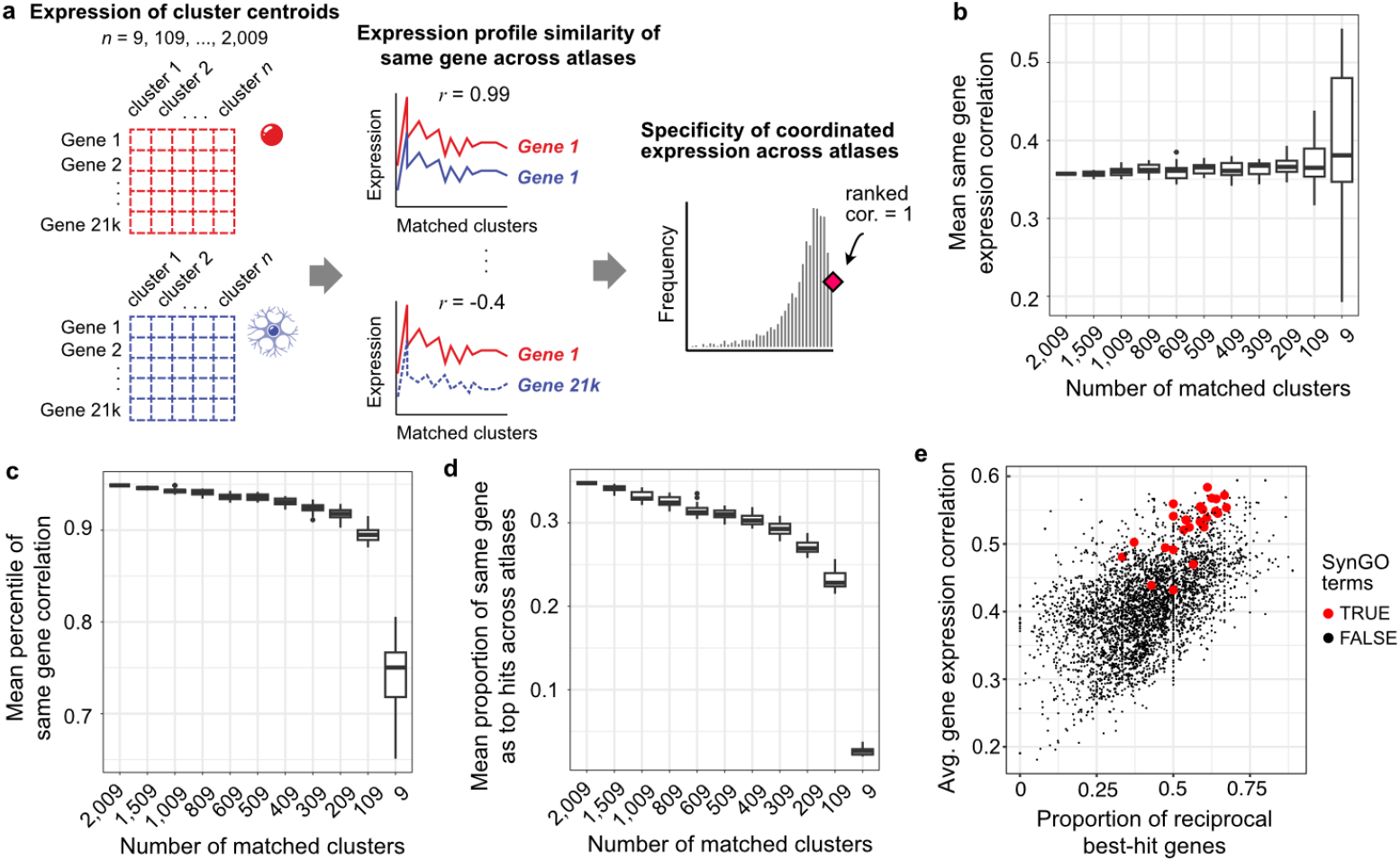
Robust and specific coordinated gene expression in reciprocally matched clusters. **a**, Schematic illustrating the calculation of coordinated expression across the centroids for a specific gene. **b**, Boxplots showing the mean Spearman correlation between matched clusters (y-axis) across increasingly smaller subsamples of the matched reciprocal clusters (x-axis). Twenty random samples were evaluated for each subsample size. **c-d**, Specificity metrics depicted through boxplots of the mean percentile of same-gene correlation and the proportion of genes with the highest same-gene correlation (y-axis) across the same cluster pairs. **e**, Scatter plot showing the relationship between the average coordinated expression across datasets and the average specificity of coordinated expression for 3,938 GO-curated gene groups. Twenty-seven SynGO-annotated gene groups are highlighted in red, indicating higher reproducibility of expression across datasets than other GO groups.

### Replicability of reciprocal clusters in additional mouse brain datasets

Next, we assessed the identification of 2,009 reciprocally matched clusters in several additional datasets. To enable fast and memory-efficient computation of cross-dataset cell type similarities, we computed pretrained models for the single-nuclei and single-cell datasets (Fig. 5a). We first applied these models to a spatial dataset that profiled expression of 109 genes in the mouse brain using the BARseq in situ sequencing technique ^17^. Of the 135 clusters identified in the BARseq study across their taxonomy, 54 showed the highest similarity to a reciprocally matched cluster using the single-cell model (odds ratio (OR): 1.10, p = 0.59), and 88 using the single-nuclei model (OR: 2.82, p < 0.0001). Combining the best hits from both models revealed 41 clusters with best hits to both sides of a reciprocally matched cluster pair. We then examined the spatial distribution of these clusters across the BARseq, MERFISH, and Slide-seq datasets. All but one pairing displayed good spatial concordance (Supplementary Table 5). Specifically, one pair exhibited a restricted pattern in the MERFISH dataset (0065 L5 IT CTX Glut_5, BARseq:L6 IT L), while three single-nucleus clusters were not detected in the Slide-seq data. Importantly, the atlases reduced annotation ambiguity, as several BARseq clusters with uncertain annotations exhibited concordant spatial patterns (Fig. 5b). Importantly, the spatial data were independent and not used in the clustering process in the original studies or our cross-dataset comparison. Despite BARseq’s limited gene panel, its spatial patterns corroborate the re-identification of the reciprocally matched cluster pairs.

**Figure 5:**
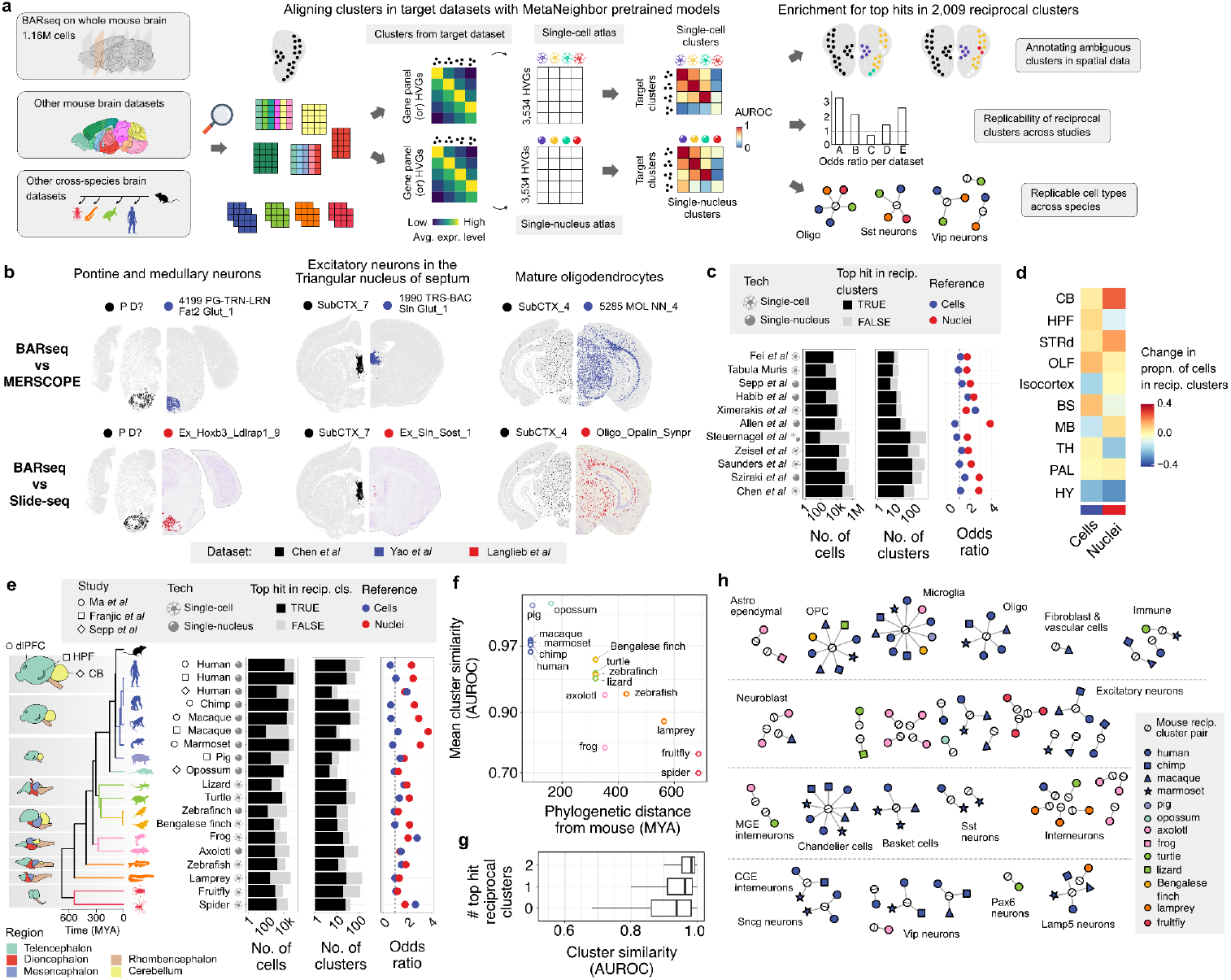
Enrichment of reciprocal matched clusters in independent spatial data and other single-cell studies. **a**, Schematic illustrating the application of a pre-trained MetaNeighbor model to the BARseq gene expression data. **b**, The held-out spatial data allows independent comparisons of expression patterns. Three BARseq (black points, left hemispheres) clusters with uninformative labels exhibit highly similar spatial patterns when compared to the top hit cluster locations in the MERFISH (blue points, top right hemispheres) and Slide-seq (red points, bottom right hemispheres) datasets. **c**, Bar plots displaying the number of cells (left) and clusters (middle) with top hit matches to either single-cell or single-nuclei reciprocal clusters (black) in different datasets. Dotplot of the odds ratios of clusters mapping to single-cell (blue diamonds), or single-nucleus (red diamonds) reciprocal clusters (right). Datasets are marked with icons to indicate the profiling method. **d**, Heatmap indicating the relative enrichment of cells in reciprocally matched clusters within aligned regions from other mouse brain datasets. Regions are listed in the same order as Figure 3f. **e**, Similar visualization as in panel c, but for brain datasets from 16 non-mouse species. Evolutionary relationships are depicted in the phylogenetic tree (left). **f**, Scatter plot showing the association between phylogenetic distance from mice (MYA: million years ago, x-axis) and the mean cluster similarity AUROC score for the top hits identified by the pre-trained models. **g**, Boxplots showing the distribution of cluster replicability AUROC scores for clusters with top hits to none, one, or both of the clusters in a reciprocal matched pair. **h**, Summary graph showing the reciprocal matched pairs as single white nodes connected to the target clusters with top hits to both sides of the reciprocal pair. Clusters in the target datasets are color-coded by species. Nodes are arranged and labelled based on broad transcriptomic cell types (ExN: excitatory neuron, InN: inhibitory neuron, MGE: medial ganglionic eminence, CGE: caudal ganglionic eminence).

To further assess replicability, we applied the pre-trained models to ten additional mouse brain datasets (Fig. 5c). These were nearly evenly split between the use of single-cell and single-nucleus sequencing. These datasets ranged from 4,000 to 385,000 sequenced cells, with the original studies clustering the cells into tens to hundreds of groups, depending on the dataset (Supplementary Table 6). The majority of cells (mean: 77.9%) and clusters (mean: 74.3%) were most similar to at least one side of a reciprocally matched cluster pair (Fig. 5c). Consistent with the BARseq findings, the top hits across datasets were enriched for the 2,009 reciprocal clusters, with stronger enrichment from the single-nucleus model (median OR: 1.91) compared to the single-cell model (median OR: 1.16). While the odds ratios were comparable between single-cell and single-nucleus preparations using the single-cell model, the single-nucleus model showed significantly higher enrichment for reciprocal matches in single-nucleus studies (median OR: 1.72 vs. 2.59, rank sum test p < 0.05). Despite the lack of fine spatial resolution in these studies, they focused on specific brain regions or used broad dissection areas. When mapped to the regions analyzed above, the cerebellum had the highest proportion of cells from replicable clusters (71%), while the hypothalamus had the lowest (21%) (Fig. 5d). A specific example is the Steuernagel et al. dataset that was focused on the hypothalamus and had the lowest proportion of cells aligned to reciprocal clusters (34%). The recurrence and enrichment of the reciprocally matched pairs across diverse datasets reinforce their utility in identifying replicable cell clusters across brain regions and sequencing techniques.

### Evolutionary conservation of reciprocal clusters

Next, we applied the pre-trained models to 19 target brain datasets from 16 species to evaluate replicability from an evolutionary perspective. Across these datasets, the number of cells and clusters was comparable to those in mouse datasets (Supplementary Table 7). On average, 74.6% of clusters which contained 74.1% of cells are most similar to at least one side of a reciprocally matched cluster pair when the model results are combined (Fig. 5e). Mirroring the mouse results, the datasets were enriched for the 2,009 reciprocal clusters, with a stronger enrichment from the single-nucleus model (median OR: 1.97) than the single-cell model (median OR: 1.06). For the single-cell model, odds ratios were higher in studies using single-cell preparations (median OR: 1.44 vs. 0.94, rank sum test, p < 0.05), while the single-nucleus model showed similar enrichment across both preparation types (median OR: 1.92 for single-cell preparations vs. 2.36 for single-nucleus studies, rank sum test, p > 0.3). The species analyzed spanned a broad evolutionary range, from humans to spiders. Notably, genetic distance from mouse was negatively correlated with mean cluster similarity scores for top hits identified by the pre-trained models (Spearman correlation = -0.89, p < 0.0001, Fig. 5f). Linking these findings to replicability, top hit scores were associated with the number of reciprocal clusters. Specifically, top hits that were not reciprocal clusters have a median AUROCs of 0.939, which increased to 0.964 for one reciprocal hit and 0.984 for clusters with reciprocal top hits from both models (Fig. 5g). Of the 1,192 cross-species target clusters, 215 had top hits corresponding to both sides of a reciprocal top hit cluster. After removing hits with inconsistent annotations at the coarse level, 104 target clusters remained (Fig. 5h). Reviewing clusters with two reciprocal top hits in four or more species highlights microglia, as well as specific clusters of oligodendrocytes, oligodendrocyte progenitor cells, Chandelier cells, and Lamp5 interneurons (Fig. 5h). Although reduced by evolutionary distance, these results suggest that the atlases and our reciprocal clusters will aid cell type identification in diverse species.

## Discussion

This study presents a comprehensive assessment of cell cluster replicability across two large-scale mouse brain transcriptomic atlases generated using single-cell and single-nucleus RNA sequencing. Our findings reveal substantial agreement between these datasets, with a significant proportion of clusters exhibiting reciprocal best-hit relationships indicative of robust cross-atlas agreement. These replicable clusters demonstrate consistent spatial localization, coordinated gene expression patterns, and are observed in independent mouse and cross-species datasets. However, our analysis also highlights the challenges associated with resolving fine-grained cellular heterogeneity and the limitations of relying solely on marker genes for cluster identification. To aid further research, we provide meta-analytic marker genes, pretrained models, and reciprocal best-hit clusters, facilitating confident application of the atlases.

Regional differences in cluster replicability, particularly the higher concordance observed in the cerebellum compared to the hypothalamus, likely arise from a combination of factors. The cerebellum’s granule cell population has limited gene expression that prevents splitting into small clusters. In addition, it has a higher proportion of nuclear transcripts due to its small cell size, may enhance its replicability across atlases that vary in subcellular sampling ^10^. In contrast, the hypothalamus’s diverse cellular composition poses a greater challenge for resolving fine-grained cell types and achieving consistent cross-atlas cluster identification. The diversity associated reduction in replicability was also observed across coronal slices. This difficulty may be further compounded by technical biases in cell isolation and sequencing processes ^18^. Such biases led to the inclusion of single-nucleus multiome sequencing data in the single-cell atlas to better capture cell populations in the midbrain and hindbrain regions ^5^. Our region-specific findings emphasize the need to account for regional factors—such as cellular heterogeneity, cell size, and technical limitations—when using transcriptomic atlases and help identify which regions require deeper sampling to improve cluster quality.

The identification of most clusters in spatial transcriptomics datasets, along with their consistent spatial distributions, aids the transition from transcriptomic clusters to defined cell types. Our findings, particularly the proximity of reciprocal best-hit cluster centroids compared to similar clusters and the repeated spatial patterns across three spatial datasets, reinforce this. Importantly, our analyses were strengthened by the independence of the spatial data from both the original clustering and our cross-atlas comparisons. However, this independence will likely be lost as spatial data is increasingly used to support the creation of transcriptomic cell-type taxonomies ^19^.

Our results align with NIH BRAIN Initiative Director John Ngai’s view that each cell cluster is a hypothesis to be tested ^20^. Unfortunately, it will be difficult to experimentally target these clusters as our analysis revealed limitations in resolving them in independent datasets. Marker genes, while effective for broadly classifying cell types, often lacked the specificity to discriminate between closely related clusters. Similarly, the “best-vs-next” AUROC scores from MetaNeighbor indicated that even reciprocal best hits sometimes struggled to distinguish between highly similar clusters. Furthermore, the average single-cell study often attains lower sampling depth for both cells and genes, further restricting the identification of fine clusters. Specifically, most of the datasets we tested the pretrained models on had a lower mean count of detected genes per cell than the atlases. This is also evident in the limited ability of the epigenetic datasets to align with the single-cell atlas. While experimentally targeting small clusters may remain challenging, integrating spatial transcriptomics offers a promising avenue for refining transcriptomic cell-type definitions.

Our findings have important implications for the development and utilization of large-scale brain transcriptomic atlases. While these resources provide invaluable insights into cellular heterogeneity, careful consideration of cluster replicability is crucial for cross-study application. Together, these whole-brain atlases provide a robust foundation for a reference brain atlas, acknowledging that some clusters may ultimately need to be merged or split. For example, both atlases identify a single, highly replicable microglia cluster that is conserved across species. In contrast, single-nucleus studies of the human brain from donors without neuropsychiatric or neurological conditions, as well as those with Alzheimer’s disease, identify multiple microglia clusters, suggesting that finer groupings may also exist in mice ^21–23^. In our analyses, many clusters do not match across atlases, possibly due to differences in subcellular and regional sampling or technical variation in sequencing methodologies. While we did align clusters with fewer than 20 cells, many small clusters may represent over-splitting due to technical artifacts or stochastic gene expression differences that do not reflect distinct cellular identities. Ultimately, improving cluster replicability will be essential for accurately characterizing cell types. Enhancing replicability will also strengthen the broader utility of transcriptomic atlases in advancing our understanding of brain organization and function.

## Methods

### Single-cell gene expression data

Single-cell sequencing data, MERFISH spatial data, and corresponding annotations were obtained from the whole mouse brain atlas by Yao et al. ^5^. The study used 317 adult C57BL/6J mice and assayed gene expression using 10X Genomics v2 and v3 kits, along with 1,687 nuclei profiled with the 10X Genomics Multiome platform. Final cell-by-gene expression count matrices were normalized using log(counts per million + 1) where necessary. Cell clusters annotated as low quality (LQ) were removed. Annotations for dissected regions were aligned to the single-nucleus atlas by manually mapping to similar or enclosing regions. For example, the hippocampus (HIP) was mapped to the hippocampal formation (HPF), while the pons (P), hindbrain (HB), and medulla (MY) were mapped to the brainstem (BS). We used Yao et al.’s marker genes and cluster taxonomy annotations for our analyses.

### Single-nucleus expression data

Single-nucleus gene expression data and annotations were obtained from the Langlieb et al. atlas ^6^. Briefly, this dataset was generated from 92 samples from 55 adult C57BL/6J mice using the 10X Genomics (v3) kit. Cluster markers were sourced from the “Whole Brain Set Cover” sheet in Supplementary Table 7 of Langlieb et al. ^6^, using only the rows where the ‘Exclude up to N Nearest Neighbors’ parameter was set to zero. Slide-seq spatial images were retrieved from the braincelldata.org website.

### Integration results from other studies

Integration results from the three companion studies in the Brain Initiative Cell Census Network 2.0 article collection were obtained from their supplementary tables. Specifically, the “Matched AIBS 10X RNA Clusters” column in Supplementary Table 4 of Liu et al. provided mappings for single-cell DNA methylome and 3D multi-omic clusters ^24^. Cluster (cl) identifiers that were no longer present in the updated Yao et al. cluster annotations were excluded from our analysis. Integration results from Songpeng et al.’s study of single-cell chromatin accessibility were retrieved from their Supplementary Table 5 ^25^. For each cluster, the one with the highest label transfer score was used, resulting in 1,040 unique single-cell clusters. To identify clusters in a transcriptomic study of spinal projecting neurons, we converted the AIBS_cluster_id values from Supplementary Table 5 of Winter et al. to cl identifiers ^26^.

### Marker-based cell calling and cluster pairing

To evaluate marker genes, we used z-scored log(CPM + 1) expression values across all cells in the atlas. For each marker set, we averaged these normalized expression values across its markers. This average marker expression was then used to rank the cells, allowing us to compute the AUROC score for all clusters in the dataset. The result was a cluster marker set by cluster AUROC matrix, with the diagonal representing the on-target AUROC scores. This matrix enabled the comparison of on-target versus best off-target AUROC scores across both atlases. Additionally, these scores were used to generate marker-based cluster pairings, where the single-nucleus cluster with the highest AUROC score for each single-cell marker set was paired. For single-nucleus clusters that were the best hits for more than one marker set, we took the pairing with the highest AUROC value.

### MetaNeighbor

We used the Python version of MetaNeighbor (https://github.com/gillislab/pyMN) to quantify cluster replicability ^14^. Default settings were used for most analyses, except for enabling the ‘run fast approximate version’ flag due to the large dataset size ^15^. Highly variable genes were selected from the 21,649 genes present in both atlases. The default symmetric AUROC matrix output was used to identify best reciprocal hits, and we also tested the asymmetric output for a more independent set of reciprocal hits.

### MERFISH spatial data

Cell-by-gene expression matrices and annotations from the Yao et al. MERFISH experiment were obtained from the Allen Brain Cell Atlas (ABC Atlas) (ID: MERFISH-C57BL6J-638850) ^5^. To assign cells to single-nucleus atlas centroids, we applied the method used by Yao et al. from the scrattch.bigcat R package. Briefly, for each cell, this method calculates the most correlated cluster centroid in the log(CPM+1) gene expression space, repeating this process 100 times after randomly subsampling the profiled genes. Cells with mean correlation values below 0.5 were assigned to clusters, in line with Yao et al.’s minimum correlation threshold. The provided Common Coordinate Framework (CCFv3) parcellation annotations were mapped to the broader dissection regions of the atlases ^27^. For example, the pons (P) and medulla (MY) were mapped to the brainstem (BS).

### Gene Ontology gene sets

Gene Ontology (GO) gene annotations dated January 2024 were obtained from the org.Mm.eg.db R package ^28^. Gene sets from the biological process aspect were filtered to have between 15 and 150 genes with coordinated expression data. The December 2023 version of SynGO was used to identify synaptic terms ^29^.

### BARseq spatial data

Single-cell expression and location data were obtained from an in situ sequencing dataset of the whole mouse brain ^17^. We analyzed the same 1,259,256 cells that remained after quality control by the original authors, grouped into 135 clusters spanning all three levels of their hierarchical taxonomy. Instead of using highly variable genes, we used all 109 genes profiled in this dataset for the MetaNeighbor pretrained models from the atlases.

### Additional single-cell brain data from mouse and other species

We assessed the utility of whole mouse brain atlases as a reference taxonomy by assessing the replicability of clusters across 11 mouse and 19 non-mouse brain datasets (see Supplementary Tables 6 and 7 for dataset details). Due to the size of the atlases, we used models pretrained on each atlas to enable fast alignment of the large number of independent datasets. These MetaNeighbor pretrained models were built using the 3,534 highly variable genes for the 5,322 single-cell and 5,030 single-nucleus clusters separatley.

Each independent dataset was filtered to include only genes present in the pretrained models. For cross-species comparisons, species-specific gene identifiers were converted to mouse orthologs using one-to-one orthology (sources provided in Supplementary Table 7). Target clusters in the independent datasets were mapped to their best-matched clusters, determined by the highest AUROC score calculated by the pretrained model. We then calculated enrichment for 2,009 reciprocal-hit clusters in the assigned top hits using Fisher’s exact test. For each target cluster in an independent dataset, we combined results from both pretrained models by counting the number of single-cell or single-nucleus best matches that are reciprocal-hit clusters. A graph visualization of the top-hit matches for cross-species target clusters was generated using the R igraph package ^30^.

Phylogenetic divergence times between species were obtained from Timetree ^31^. Species silhouettes were sourced from www.phylopic.org (public domain), and brain illustrations were adapted from Lamanna et al. ^32^.

## Data Availability

The pretrained MetaNeighbor models, AUROC matrices, and markers are publicly available on Figshare (https://figshare.com/articles/dataset/Files_for_Cluster_replicability_in_single-cell_and_single-nucleus_atlases_of_the_mouse_brain/28462349).

## Code availability

Scripts used in this study are publicly accessible on GitHub at https://github.com/gillislab/mouse_brain_cluster_replicability/.

## Author Contributions

J.G. conceived and supervised the project. L.F. and H.S. performed analyses. L.F. drafted the manuscript. All authors interpreted the results and assisted in writing and revising the paper.

## Acknowledgements

L.F. received support from R01MH133181 and J.G. from both R01MH133181 and R01MH113005. The content is solely the responsibility of the authors and does not necessarily represent the official views of the National Institutes of Health.

## Supplementary Tables

Supplementary Table 1 - Identifiers, cluster sizes, and AUROC scores for the 2009 reciprocal best hit pairs.

Supplementary Table 2 - Enrichment of the 2009 reciprocal best hits in the 34 cell classes in the single-cell atlas.

Supplementary Table 3 - Enrichment of the 2009 reciprocal best hits in the 161 single-nuclei metaclusters.

Supplementary Table 4 - Average correlation and proportion of best matched gene-gene correlations from the coordinated expression analyses for GO groups.

Supplementary Table 5 - Manual curation results and characteristics of the 41 BARseq clusters with top hits to reciprocal cluster pairs.

Supplementary Table 6 - Information for additional mouse brain datasets re-analysed. Supplementary Table 7 - Information for cross-species brain datasets re-analysed.

Supplementary Table 8 - Identifiers, cluster sizes, and AUROC scores for the 612 high confidence reciprocal best hit pairs.

## Supplementary Figures

Supplementary Figure 1: Number of reciprocal best-hit clusters achieving a best-versus-next AUROC of 0.95 or greater (y-axis) for varying numbers of MetaMarkers (x-axis) for single-cell (blue) and single-nuclei (red) datasets.

Supplementary Figure 2: Smoothed trends of discrimination performance (negative log_10_ Mann–Whitney U test p-value) versus cluster size (log-scaled) for best-versus-next cluster comparisons. The estimated trend lines were computed using Generalized Additive Models (GAMs) via geom_smooth() in R’s ggplot2, with shaded areas indicating 95% confidence intervals. Solid lines depict the association within a dataset and dashed lines show it across datasets for reference clusters derived from cells (blue) and nuclei (red).

