## Supplementary figures and images for "Cluster replicability in single-cell and single-nucleus atlases of the mouse brain"

### Supplementary Figure 1

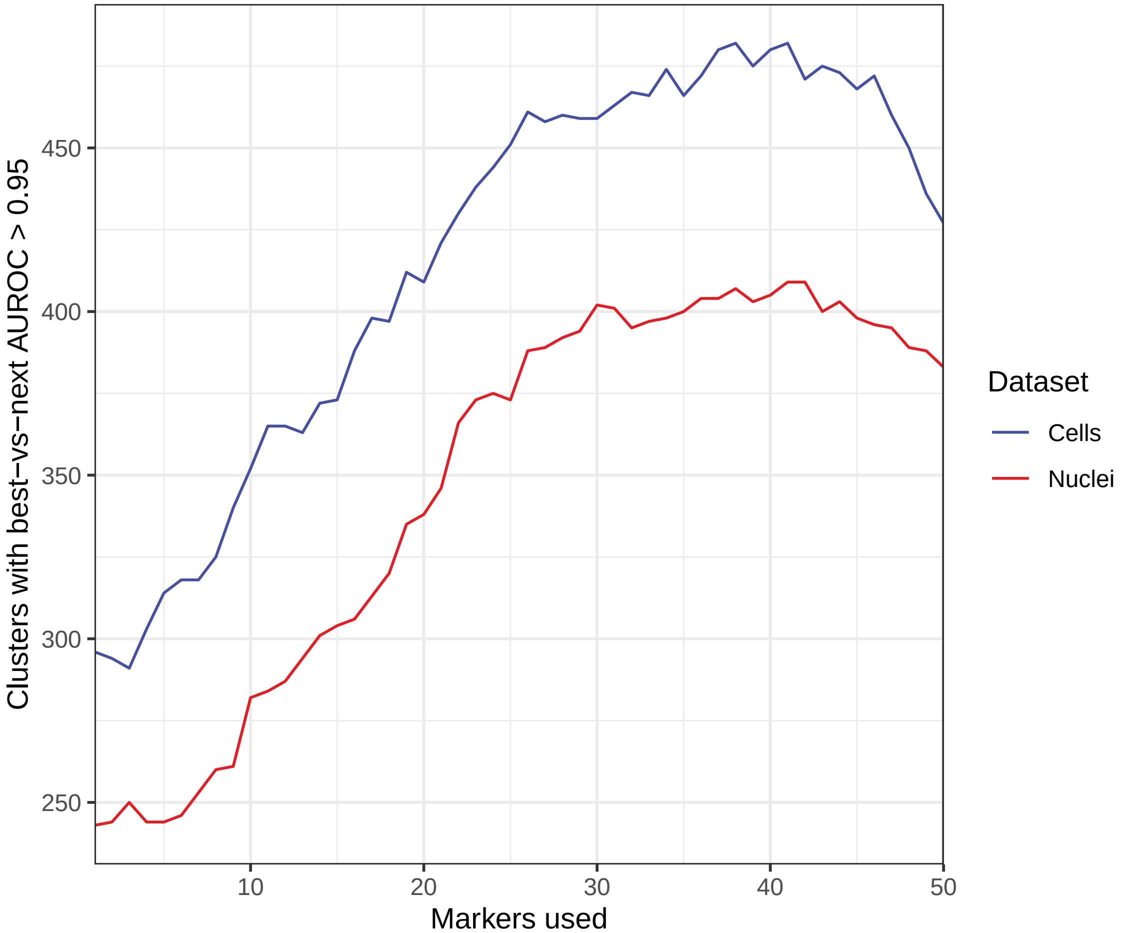

### Supplementary Figure 2

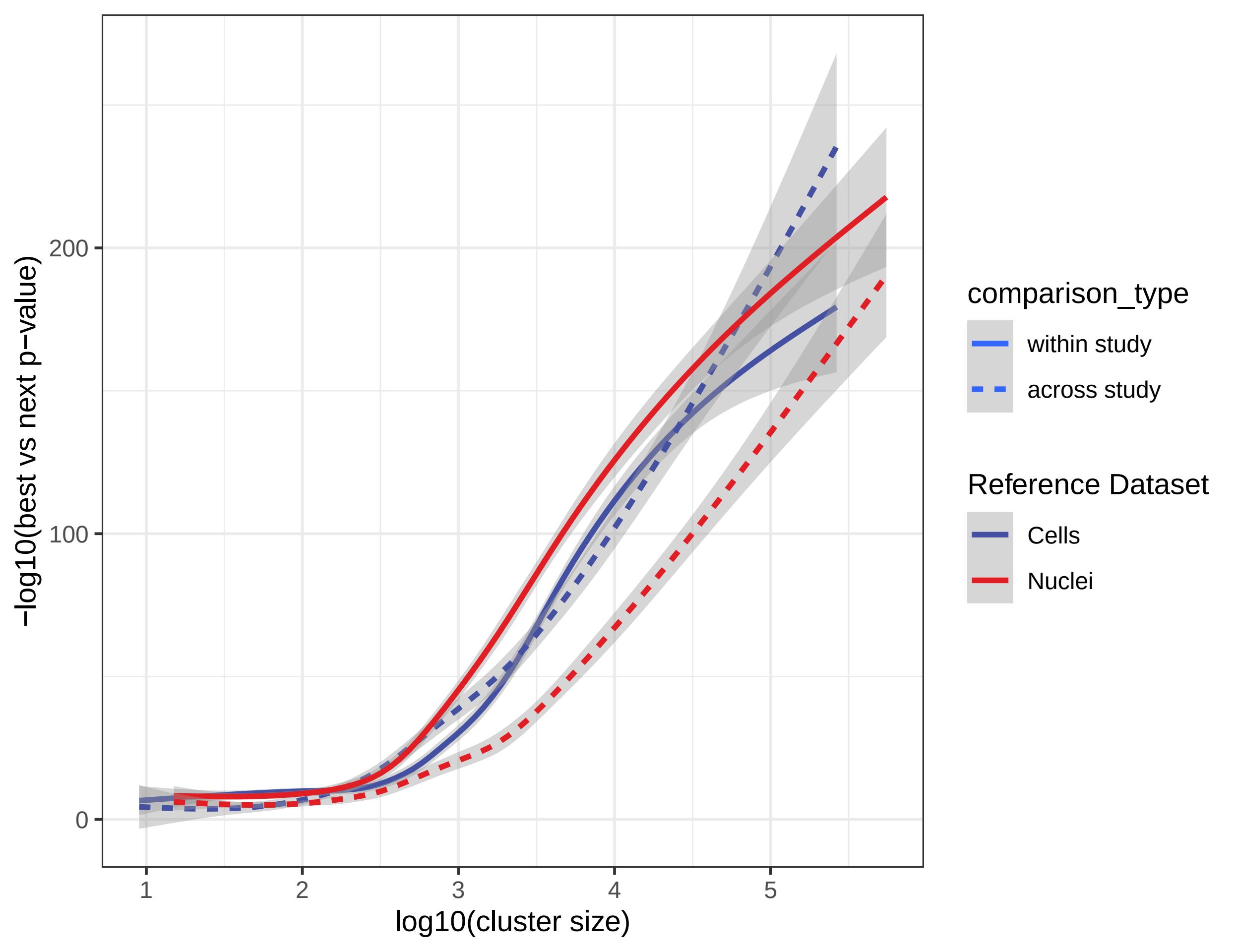
