## Supplementary Notes for "Cluster replicability in single-cell and single-nucleus atlases of the mouse brain"

### Supplementary Note 1: Metamarker identification

Next, we revisited marker gene testing, as our initial analysis showed that marker genes lacked the specificity to distinguish similar clusters. Using MetaMarkers with default parameters, we identified robust markers for the 2,009 reciprocal cluster pairs [1]. Since this method compares the target cluster to all other cells, we evaluated marker performance in a local context. Using the top four markers, the mean best-vs-next AUROC reached 0.697 for the single-cell data and 0.664 for the single-nucleus data. Applying a stricter threshold, only 244 nuclei clusters and 303 cell clusters could be locally discriminated with an AUROC greater than 0.95 (Supplementary Figure 2). The maximum discrimination was reached using the aggregate expression of 41 markers, with 409 single-nucleus clusters and 482 single-cell clusters passing the 0.95 threshold. Overall, this suggests that small sets of marker genes have limited ability to distinguish thousands of transcriptomically identified clusters accurately.

### Supplementary Note 2: Refining cluster alignment

To increase the replicability of reciprocal matches between the atlases, we applied stricter thresholds to generate a set of high-confidence cluster pairs. Instead of selecting reciprocal pairs based on the symmetric similarity matrix, where reference and target cluster AUROC scores are averaged, we used the asymmetric version and filtered out pairs with AUROC scores below 0.99. Additionally, we required a best-vs-next AUROC score greater than 0.6 to ensure adequate discrimination among similar clusters. The remaining 612 reciprocal best hit pairs include 14% of the single-cell and 29.4% of the single-nuclei cells (Supplementary Table 8). Compared to the original 2,009 pairs, these 612 showed improved performance across several metrics. For instance, the correlation between paired cluster sizes increased from 0.51 to 0.59. Spatially, there was stronger agreement, with 86.1% of the pairs matching in the top dissection region, compared to 80.8% for the 2,009 pairs. The mean centroid distance in the MERFISH data also decreased, from 0.428 mm to 0.34 mm. Additionally, the coordinated expression metrics remained consistent between the two sets. Despite these improvements, the substantial reduction in the number of paired clusters prompted us to focus primarily on the 2,009 reciprocal pairs, as the differences in our assessments were marginal.

To improve alignment between the two atlases, we tested several strategies for merging clusters. For example, if two clusters showed high similarity relative to a reference cluster in the other atlas (0.4 < best-vs-next AUROC < 0.6), we merged them. We also explored merges guided by the provided cell type taxonomies. While these methods reduced the number of unpaired clusters, they did not significantly increase the number of reciprocal top-hits beyond 2,009.
